# Vamos en bici: Study protocol of an investigation of cognitive and neural changes following language training, physical exercise training, or a combination of both

**DOI:** 10.1101/2022.01.30.478181

**Authors:** Elisabeth Wenger, Sandra Düzel, Sarah E. Polk, Nils C. Bodammer, Carola Misgeld, Johanna Porst, Bernd Wolfarth, Simone Kühn, Ulman Lindenberger

## Abstract

As the relative number of elderly people as well as the average life expectancy increases, identifying potential means to alter the ongoing trajectory of aging and specifically the trajectory of cognitive performance is of great importance. Some modifiable lifestyle factors, such as physical and cognitive activity, have shown positive effects on cognition and brain structure, and the combination of the two might even show a boosted, interactive effect. In this study protocol, we describe in detail our data which was acquired to explore how cognitive stimulation in the form of acquiring a new language, physical exercise on stationary bikes or the combination of the two interventions affect brain structure, cognitive performance, and psychosocial functioning. One-hundred and forty-two older healthy adults (63–78 years) were randomly assigned to one of four six-month intervention programs, comprising *(a)* foreign language learning, *(b)* physical exercise training on a bicycle ergometer, *(c)* a combination of language learning and physical exercise, or *(d)* a book club (serving as an active control condition). We collected a rich neuroimaging data set, comprising T_1_-weighted structural, resting state functional, high resolution hippocampal, myelin water fraction, diffusion-weighted, arterial spin labeling, and multi-parameter images. Using a cognitive battery, we collected data from the domains of episodic memory, working memory, perceptual speed, and fluid intelligence. We performed comprehensive physical assessments including cardiopulmonary exercise testing, and additionally collected data on psychosocial functioning (e.g., well-being, perceived stress, control beliefs). We assume that physical activity boosts brain plasticity *per se* by inducing structural and neurochemical changes in brain regions that are important for learning and memory and therefore may facilitate the effects of cognitive training. (269 words)

## Introduction

Living a long, healthy life — this is what everyone is striving for. The very good news is that life expectancy has been steadily increasing ever since 1840 (Vaupel et al., 2003). While back in 1960, average life expectancy was still around 60 years, in 2020 it was already over 80. Around the world, and especially in Europe, the older segment of the adult population is therefore steadily growing in size and proportion. In dealing with the demographic change, it is important what people make out of the added years of their lives.

With increasing age, even healthy and able individuals experience some decline of cognitive performance (Lindenberger, 2014). Many older adults feel that their memory does not serve them as well as when they were younger. Not all forms of human memory are equally affected by advancing age: positive age gradients have been found for semantic memory, while episodic memory is considered to be the form of long-term memory that displays the largest degree of age-related decline (Rönnlund et al., 2005). Longitudinal studies that apply appropriate control for practice effects (i.e., test-retest effects) indicate that episodic memory performance remains relatively stable until about 60–65 years of age, after which accelerating decline is typically observed (Nyberg et al., 2012).

As the relative number of elderly people as well as the average life expectancy is increasing, identifying potential means to change the ongoing trajectory of aging and specifically the trajectory of episodic memory is of great importance (Hertzog et al., 2009; Lindenberger, 2014). Leading an intellectually challenging, physically active, and socially engaged life may be the best guess we currently have to provide optimal grounds for enhanced cognitive stability and growth and potentially even postponing cognitive decline at least to some extent (Lindenberger, 2014). Both cognitive as well as physical training have been shown to occasionally improve cognitive performance of healthy adults across the lifespan, even in old age, drawing on the brain’s lifelong plasticity (Lövdén et al., 2013).

Indeed, there are different studies demonstrating positive effects of various kinds of cognitive activity on brain structure (for example, studying for a medical exam, years of training to become a taxi driver, performing a spatial navigation training, or intensely studying a foreign language) (Draganski et al., 2006; Lövdén et al., 2012; Mårtensson et al., 2012; Woollett & Maguire, 2011). There are even more studies showing positive effects of physical exercise on brain structure and cognition (Ahlskog et al., 2011; Erickson et al., 2011; Kleemeyer et al., 2015; Maass et al., 2015; Macpherson et al., 2017; Ngandu et al., 2015; Prakash et al., 2015). Crucially, it has been proposed that physical and cognitive exercise might interact to induce even larger functional benefits (Bamidis et al., 2014; Hötting & Röder, 2013; Kempermann et al., 2010; Kraft, 2012; Lustig et al., 2009). Several intervention studies are in line with this hypothesis, demonstrating larger benefits on cognitive test performance for combined physical and cognitive activity than for each activity alone. Those studies comprise observational cross-sectional (Eskes et al., 2010), longitudinal (Karp et al., 2006), as well as controlled interventional designs (Fabre et al., 2002; Oswald et al., 2006). From an evolutionary perspective, physical and cognitive challenges were highly interwoven (Kempermann et al., 2010). It is therefore reasonable to assume that the neurobiological mechanisms triggered by physical and cognitive exercise go hand in hand. Physical exercise might increase the potential for neurogenesis and synaptogenesis whilst cognitive exercise guides it to induce positive plastic change (Fissler et al., 2013). It has been observed in animal work, that running combined with environmental enrichment leads to the most pronounced increase in the number of new neurons in dentate gyrus compared to either training alone, thus demonstrating an additive effect of exercise and enrichment (Fabel et al., 2009). However, it also needs to be noted that a number of studies found no additional or synergistic effects (Barnes et al., 2003; Legault et al., 2011; Shatil, 2013).

To summarize, accumulating and promising evidence suggests that plastic changes are less pronounced but possible in old age, and might be enhanced by combining both physical and cognitive activities. In this study, we therefore make use of two powerful variants of modifiable lifestyle factors and explore how interventions deliberately embedded into participants’ daily lives, namely *(a)* cognitive stimulation in the context of acquiring a new language, *(b)* physical exercise and *(c)* the combination of the two interventions affect brain structure and function, cognitive performance, and psychosocial functioning. We assume that physical activity facilitates brain plasticity *per se* by inducing structural and neurochemical changes in brain regions that are important for learning and memory and therefore may boost the effects of cognitive training.

## Methods

### Participants and study design

Volunteers (*N* = 201) recruited through newspaper advertisements and previous participation in studies^1^ were included if they met the following inclusion criteria: no MRI contraindications; able to meet the time requirements of the study (ca. 45 minutes a day for six months); right-handed; between 63 and 78 years old at the start of the study^2^; no history of head injury, medical (e.g., heart attack, cancer), neurological (e.g., epilepsy), or psychological disorder (e.g., depression); no prescription of medication affecting memory function. To ensure similar baseline levels of language and exercise, participants were excluded if they could speak a romance language or were proficient in more than one language besides German, or if they already engaged in aerobic exercise more than once every two weeks.

Qualifying volunteers were pseudo-randomly assigned to one of four groups: (1) an active control group that participated in a book club (ACG), (2) a language training group (LG), (3) an exercise training group (EG), (4) a combined language and exercise training group (L+EG), and were first invited for group-specific information sessions at the Max Planck Institute for Human Development, Berlin, where the general set-up of the study was explained and details about the time commitment and the planned assessments were given. Participants were unaware of the other existing groups, respectively. They were then invited to a physical assessment including cardiopulmonary exercise testing, where 22 participants were excluded due to pre-existing medical conditions. Finally, participants underwent an initial magnetic resonance imaging (MRI) session and completed a comprehensive cognitive battery before beginning their assigned training (pre-intervention, T1). Nineteen participants dropped out before the training started due to disinterest and one additional participant was excluded due to claustrophobia. After three months of training (mid-intervention, T2), MRI and cognitive sessions were repeated, consisting of the same tests as at T1. After six months (post-intervention, T3), MRI and cognitive measures were acquired again, and participants underwent a physical assessment. During the intervention, 17 participants dropped out due to health problems, disinterest, time constraints, or unspecified reasons, leaving a total of 142 participants who completed the study. *A posteriori*, participants were considered fully compliant if they completed at least 1890 minutes (equaling approximately 90 minutes per week for 21 weeks) of study-related activity during the whole intervention. Compliance was assumed to be unrelated to outcomes of interest at baseline (e.g., hippocampal subfield volume, episodic memory) but may have affected the rate or magnitude of change in these outcomes over time, so non-compliant participants should be included in an analysis at baseline only. Please see Figure 1 for a detailed flow-chart of the participant recruitment process and an overview of the study design.

**Figure 1.**
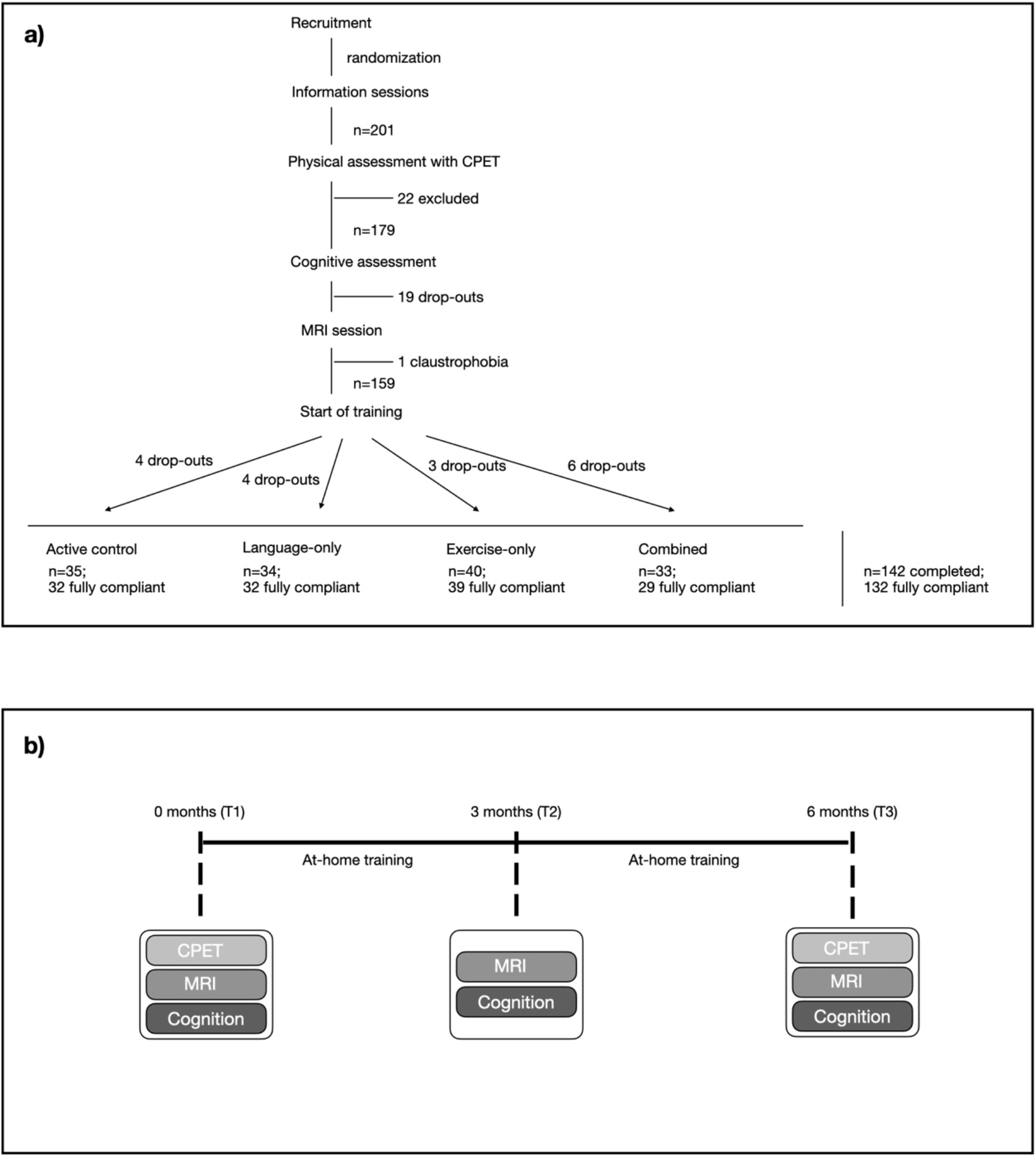
Overview of recruitment process and study design. While 159 participants started with the intervention, 17 dropped out during the intervention due to health-related issues or time-constraints, leaving 142 participants who completed the full study. Of these 142, 10 participants invested less than 1890 min of training over the whole study period and were therefore considered as non-compliant. The final sample of fully compliant participants therefore comprises 132 participants.

The ethics committee of the German Society for Psychology (DGPs) approved the study and written informed consent was collected from all participants.

### Interventions

#### Active control group

Participants in the active control group (ACG; *n* = 35) were instructed to read pre-selected German literature on a tablet for 45 minutes daily for at least six days per week. These participants attended weekly group sessions during which they discussed literary excerpts led by external “facilitators” (http://shared-reading.de/). Three participants did not fully comply with the intervention conditions, thus, their data should only be included in analyses focusing on baseline data only.

#### Language group

Participants in the cognitive training group (LG; *n* = 34) learned Spanish using the Babbel application on a tablet (Lenovo TB2-X30L TAB) for about 30 minutes a day (or until they completed one lesson) for six days a week. Participants were also instructed to chronologically read pre-selected literature on the tablet for an additional 15 minutes a day, amounting to about 45 minutes of study-related activity each day. They also participated in weekly one-hour group sessions (5–10 participants per session) at the institute consisting of a Spanish class led by an external instructor. Two participants in the language group did not fully comply with the intervention conditions and should be used in baseline analysis only.

#### Exercise group

Participants in the exercise group (EG; *n* = 40) engaged in moderate at-home exercise three to four times per week using a bicycle ergometer (DKN Ergometer AM-50) and a personalized interval training regime programmed onto a tablet, which were synced via Bluetooth. The training initially lasted 30 minutes at an individually defined intensity, and the difficulty increased by three minutes per interval and by three to four Watts (W) approximately every two weeks. After completing a training session, participants could indicate if they found the training too easy or difficult, and the intensity could be remotely adjusted accordingly. In this way, the training was highly personalized so that participants would not be discouraged by a difficult exercise program. Participants in the exercise group were also instructed to read pre-selected literature on the tablet for an additional 15 minutes on days when they trained and for 45 minutes on the other days, again leading to a mean of 45 minutes of study-related activities per day. Finally, those in the exercise group participated in weekly one-hour group sessions at the institute consisting of toning and stretching, led by an external instructor. One participant did not fully comply and one experienced prohibitive technical difficulties during the intervention period; again this should only be included in baseline analyses.

#### Combined language and exercise group

Participants in the combined language and exercise training group (L+EG; *n* = 33) both learned Spanish using Babbel six days per week and engaged in the interval training on an ergometer for three to four days per week, as described above. Participants were completely free to decide at what time of the day to do the training, what to do first, or how much time they let pass between the two training elements. Doing approximately 30 min of language training 6 times a week, and 30 min of exercise training on 3-4 days a week led to a mean of 45 minutes of study-related activities per day, just like in the other 3 groups. They also joined weekly Spanish classes (separated from the cognitive training group) at the institute in groups. Four participants did not fully comply with intervention conditions and should only be used in baseline analyses.

### Data acquisition

Measurements included two physical assessments including cardiopulmonary exercise testing (pre- and post-intervention), three MRI scans, and three cognitive sessions (pre-, mid-, post-intervention; see Figure 1b). Data were also collected immediately before, during, and immediately after at-home study-related activity via the tablets and ergometers (from the exercise group), as well as whenever participants wore an activity tracking device (Fitbit watch). Further details are described below.

### Comprehensive physical assessments with cardiopulmonary exercise testing

The physical assessments conducted at Charité Berlin under medical supervision provided various measures of individual fitness levels pre- and post-intervention, as well as a baseline for the personalized training programs prepared for participants in the EG and L+EG. The appointments included comprehensive anamnesis, a physical check-up with anthropometric and cardiovascular measures, evaluation of cardiopulmonary and vascular function using bicycle ergospirometry, and blood sampling.

#### Physical check-up

Body weight in kilograms and height in meters were measured and body mass index (BMI = kg/m²; using the Seca 285 DP measuring station, Seca GmbH, Hamburg, Deutschland) was calculated. Waist and hip circumference were measured and waist-to-hip ratio (WHR) (WHO Expert Consultation, 2008) was calculated. WHR is a simple measure of body fat distribution ratio, with higher values indicating greater cardiovascular health risk due to greater amounts of visceral abdominal fat (WHO Expert Consultation, 2008). Body fat was measured using both the seven-site (Jackson & Pollock, 1978) and ten-site skinfold measurement technique (Johnsen & Scholz, 1989) using calipers. Resting heart rate was measured using a 12-channel resting electrocardiogram (Cardio 200 Suction ECG, Custo med GmbH), which was also used to ensure that participants were healthy enough to undergo ergospirometry, and auscultatory measurements of systolic and diastolic blood pressure (BP) were taken from each arm.

#### Blood sampling

Indicators of general health and metabolism as well as markers of neuronal plasticity were acquired via blood sampling. Three 5 ml venous blood samples were drawn in additive-free vacuum containers (EDTA 7.5 ml vial; heparin tube 7.5 ml; S-Monovette® 7.5 ml Z-Gel, Sarstedt) immediately before assessing performance diagnostics (described below). Vacuum containers were inverted 10 times after being drawn and left upright to clot for at least 30 minutes in a refrigerator at 4 ºC. One sample was then centrifuged for 15 minutes at 1100 g at 4 ºC, and 100 µl aliquots of serum were transferred to three 100 µl Eppendorf Safe-Lock tubes and one 1.8 ml Thermo tube (Thermo Fischer Scientific); the remaining serum was transferred to a 2 ml Eppendorf Safe-Lock tube. Serum samples were stored at –80 ºC until being transported to the laboratory. Another 5 ml venous blood sample (S-Monovette® 7.5 ml Z-Gel, Sarstedt) was drawn immediately following performance diagnostics, which was also centrifuged as described above. Serum from this sample was aliquoted into three 100 µl Safe-Lock tubes (100 µl of serum each) and stored at –80 ºC until being transported with the other blood samples.

The 5 ml venous blood samples in the EDTA and heparin tubes were sent to Labor28 (https://web.labor28.de/) for analysis. Parameters of general health and metabolism were assessed, including parameters indexing liver health: high- and low-density lipoprotein (HDL and LDL), C-reactive protein (CrP), triglycerides; and parameters of kidney health, namely urea and creatinine. A hemogram or complete blood count was also performed to assess levels of monocytes, lymphocytes, leukocytes, and neutrophils. Additional parameters targeting neuroinflammation included interleukin 1 beta (IL-1ß), interleukin 6 (IL-6), interleukin 10 (IL-10), interleukin-1 receptor antagonist (IL-1RA), interleukin 18 (IL-18, also known as interferon-gamma inducing factor), tumor necrosis factor (TNF), human soluble tumor necrosis factor receptor antagonist (sTNF-RA), cortisol, dehydroepiandrosterone (DHEA), and cytomegalovirus antibodies (CMV).

The aliquoted serum samples were sent to the University Hospital Würzburg to quantify levels of brain-derived neurotrophic factor (BDNF), insulin-like growth factor 1 (IGF-1), insulin-like growth factor-binding protein 3 (IGFBP-3), and cortisol in blood serum immediately before and after physical exertion.

Another batch of aliquoted serum samples was sent to University Clinic Hamburg-Eppendorf to quantify levels of glycosylphosphatidylinositol (GPI)–specific phospholipase D1 (Gpld1), a GPI-degrading enzyme derived from liver, that has been found to increase after exercise and to correlate with improved cognitive function (Horowitz et al., 2020).

#### Performance diagnostics

To assess performance diagnostics, participants were instructed to pedal at a constant rate of 60–70 RPM on a bicycle ergometer during the entire protocol, which consisted of a three-minute rest phase, an exertion phase with a starting resistance of 20 W, which increased by 20 W every three minutes until participants reported they had reached maximum exertion, and a five-minute recovery phase with no resistance (as defined by (Bosquet et al., 2002; Braumann et al., 2004). During this protocol, ergospirometry was performed, cardiac activity was constantly monitored, and lactate levels were measured stepwise via earlobe blood sampling, as described in detail below.

##### Borg Rating of Perceived Exertion (RPE) Scale

After the end of each interval, participants reported their subjective level of exertion using the Borg Scale (Borg, 1982). This is a 15-point scale used for the evaluation of physical activity intensity level. Participants rate their total feeling of exertion from 6 (“no exertion at all”) to 20 (“maximum exertion”).

##### Lactate

Lactate kinetics recording is a standard measurement method for assessing endurance performance. Lactate levels were measured during the last 10 seconds of each interval by sampling 20 µl of capillary blood from the earlobe, which was mixed with 1 ml of a hemolysis solution. A lactate curve was calculated using lactate level from at least five intervals, heart rate, blood pressure, and Borg Scale scores. This lactate curve was then used to estimate the lactate threshold, which is the point at which lactate starts accumulating in the muscles (Brooks, 1985). Additionally, we also calculated the individual anaerobic threshold following Dickhuth and colleagues (1991).

##### Cardiac activity

At the end of each 20 W interval, cardiac activity was recorded with the Cardio 110 BT (Custo med GMbH) and the Custo diagnostic software (Version 4.4.4 [x64]).

##### Blood pressure

Blood pressure was measured 40 seconds before the end of each interval using the Riva-Rocci/Korotkoff method, in which an upper arm cuff is applied and a stethoscope was used to listen to the Korotkoff sounds.

##### Ergospirometry

Ergospirometry, along with lactate kinetics, is the current gold standard for determining endurance performance (Scharhag-Rosenberger & Schommer, 2013). This non-invasive method is suitable for measuring the performance diagnostics of both athletes and untrained individuals. Notably, older participants may not reach their absolute maximum achievable workload, but rather a subjective maximum. Thus, in the following, we refer to the final measurements as measurements at peak, which may not necessarily reflect participants’ absolute maximum.

Respiration and gas metabolism were measured using the Quark Cardio Pulmonary Exercise Test (CPET; COSMED) with the standard Breath-by-Breath setup, and the V2 Mask (Hans Rudolph, Inc.), which is a mask that covers the mouth and nose, and is fastened to the back of the head. Parameters were recorded with the OMNIA software (Version 1.6.2, COSMED), including peak oxygen uptake (VO_2_peak) measured in 30-second intervals, relativized by body weight in kilograms (Röcker, 2010), which is a gold standard measurement of endurance performance, as well as respiratory quotient and ventilation (at peak and at the ventilatory threshold), power (stepwise), and heart rate (constant). Additionally, using the lactate measurements, heart rate and power at the lactate threshold as well as the ventilatory threshold and peak were calculated.

#### Magnetic resonance imaging

Participants were scanned at pre-, mid-, and post-intervention. MR images were acquired using a Siemens Tim Trio 3T MR scanner (Siemens Medical Systems, Erlangen, Germany; VB17a software version) using a standard radio-frequency (RF) 32-channel receive head coil and RF transmit body coil. Acquisition details are described below.

##### T_1_-weighted MPRAGE

Structural images were obtained using a three-dimensional T_1_-weighted magnetization-prepared gradient-echo sequence (MPRAGE) (TR = 2500 ms, TE = 4.77 ms, TI = 1100 ms, field of view (FOV) = 256 × 256 × 192 mm^3^, flip angle = 7º, bandwidth = 140 Hz/pixel, isometric voxel size = 1 mm^3^, total time of acquisition (TA): 9:20 min).

##### Resting state acquisition

We used a T_2_*-weighted multiband-EPI sequence of the Center of Magnetic Resonance Research (CMRR, Minneapolis) of 8 minutes length, sensitive to Blood Oxygenation Level Dependent (BOLD) contrast (TR = 2000 ms, TE = 30 ms, FOV = 216 × 216 × 129 mm^3^, flip angle = 80°, slice thickness 3.0 mm, distance factor = 20%, in-plane resolution = 3 × 3 mm^2^, 36 axial slices, using GRAPPA acceleration factor 2, phase encoding direction anterio-posterior, TA = 8:08) to acquire whole brain functional images for resting state analyses. Slices were acquired in an interleaved fashion, aligned to the line connecting the lower borders of genu and splenium of the corpus callosum. For the purpose of correcting for distortions due to B_0_ inhomogeneities, we also acquired ten image volumes with inverted phase-encoding direction (TA = 0:28).

##### High-resolution hippocampal scans

A high-resolution, T_2_-weighted 2D turbo spin echo (TSE) sequence was acquired, perpendicularly oriented to the long axis of the right hippocampus, with in-plane resolution = 0.4 × 0.4 mm^2^, slice thickness = 2.0 mm, 31 coronal slices, image matrix 384 × 384, with TR = 8510 ms, TE = 50 ms, flip angle = 120°, turbo factor 15 applying hyperechoes, bandwidth 99 Hz/pixel, 1 average per acquisition, TA = 7:13 min.

##### Multi-echo GRASE sequence for myelin water fraction mapping

The T_2_ relaxation images were acquired using a 3D multi-echo (ME) gradient and spin echo (GRASE) sequence, which was generously contributed by Dr. Jongho Lee (Seoul National University) and shared by Dr. Jeffrey A. Stanley (Wayne State University), with the following parameters: TR = 1000 ms, number of echoes (TEs) = 32, first TE = 10 ms, inter-echo spacing = 10 ms, EPI factor = 3, FOV = 240 × 180 × 128 mm, in-plane resolution = 1.5 × 1.5 mm^2^, slice thickness = 4 mm, slice partial Fourier 6/8, no parallel imaging, TA = 16:03.

##### Diffusion-weighted imaging

Two-shell diffusion-weighted images were obtained with a single-shot diffusion-weighted spin-echo-refocused echo-planar imaging sequence with the following parameters: TR = 9700 ms; TE =120 ms; 62 slices; FOV = 224 × 224 mm; slice thickness 2.0 mm; *b*-values 2850/710 s/mm^2^; 60/30 directions; with 10 non diffusion-weighted images distributed equidistantly throughout; GRAPPA acceleration factor = 2; 2 mm isotropic voxels; phase-encoding direction anterior-posterior; acquisition time = 16:41 min. Five additional images inverting the phase-encoding direction were acquired without diffusion weighting; TA = 1:29 min.

##### FLAIR sequence for WM hyperintensities

We collected a fluid attenuation inversion recovery (FLAIR) image volume in order to quantify white matter hyperintensities and to aid visual inspection of abnormalities if any were detected. The following parameters were used: TR = 10000 ms, TE = 97 ms, flip angle = 125°, TI = 2601.3 ms, voxel size = 1 × 1 × 3 mm, TA = 3:42 min.

##### Arterial-spin labeling (ASL)

A multi-TI, background-suppressed pulsed ASL (PASL) sequence with a flow alternating inversion recovery (FAIR) labeling and Q2TIPS saturation pulses, contributed by Dr. Matthias Günther (MEVIS, Bremen; Günther et al., 2005) and implemented with the help of Dr. Federico Samson-Himmelstjerna, was used with the following parameters (Martin et al., 2015): 14 TIs (start of the time series at 100 ms, increment 250 ms, end at 3350 ms), with a bolus length of 1800 ms, and two background suppression pulses were acquired. Readout of two averages was performed via 3D-GRASE with the following parameters: TR = 3800 ms, TE = 17.24 ms, flip angle (refocusing pulses) = 100°, EPI factor 24, turbo factor 20, number of segments = 2, FOV = 256 × 192 × 110 mm^3^, voxel size = 4.0 × 4.0 × 5.5 mm^3^, bandwidth 2298 Hz/Px, TA = 7:07. Additionally, we acquired 2 M_0_ images with R > L and L > R phase-encoding directions (TA = 50 s each) in order to correct geometric distortions in the CBF-maps.

##### Multi-parameter mapping (MPM)

The MPM protocol was generously provided by Dr. Nikolaus Weiskopf and Dr. Martina Callaghan (Weiskopf et al., 2021) and comprised one GRE-field map quantifying the inhomogeneities of the static magnetic field (B_0_), one RF transmit field map (B_1^+^_), and three multi-echo 3D FLASH (fast low angle shot) sequences. The B_0_ gradient echo field mapping sequence was acquired with the following parameters: 64 transverse slices, slice thickness = 2 mm with 50% distance factor, TR = 1020 ms, TE TE1/TE2 = 10/12.46 ms, flip angle a = 90°, matrix = 64 × 64, field of view (FOV) = 192 × 192 mm, right-left phase encoding (PE) direction, bandwidth (BW) = 260 Hz/Px, flow compensation, TA = 2:14 min.

Maps of the local RF transmit (B_1_ ^+^) field were estimated from a 3D EPI acquisition of spin and stimulated echoes (SE and STE) with different flip angles. The following parameters were used: 4 mm isotropic resolution, matrix = 64 × 48 × 48, FOV = 256 × 192 × 192 mm, parallel imaging using GRAPPA factor 2 × 2 in PE and partition directions, TR = 500 ms, TE_SE/STE_/mixing time = 39.06 ms/33.80 ms. Eleven pairs of SE/STE image volumes were measured successively employing decreasing flip angles (applied in an α–2α–α series) from 115° to 65° in steps of -5° (Akoka et al., 1993; Lutti et al., 2010). TA was 3 min.

The three different multi-echo FLASH sequences were acquired with predominantly T_1_ weighting (T_1_w), proton density weighting (PDw), or magnetization transfer weighting (MTw) by appropriate choice of repetition time (TR) and flip angle α (T_1_w: TR/α = 24.5 ms/21°; PDw and MTw: TR/α = 24.5 ms/6°) and by applying an off-resonance Gaussian-shaped RF pulse (4 ms duration, 220° nominal flip angle, 2 kHz frequency offset from water resonance) prior to excitation in case of the MTw sequence version. Multiple gradient echoes with alternating readout polarity were acquired at six equidistant echo times (TE) between 2.34 ms and 14.04 ms for the T_1_w and MTw acquisitions with two additional echoes at TE = 16.38 ms and 18.72 ms for the PDw acquisition. Additional parameters were: readout bandwidth = 465 Hz/pixel, GRAPPA parallel imaging with an acceleration factor of 2 in the phase-encoding (anterior-posterior) direction (outer/slow phase encoding loop), 6/8 partial Fourier acquisitions in the partition (left-right) direction (inner/fast phase encoding loop), 1 mm isotropic resolution, 176 slices per slab, FOV = 256 × 240 mm, TA of each of the three FLASH sequences = 7:03 min.

#### Cognition

Participants completed a comprehensive battery of cognitive tests pre-, mid-, and post-intervention (see Figure 1b for a study design overview) that was assessed within a group testing session of 4 to 4.5 hours (including extensive breaks with refreshments and snacks). The tasks pertained to various cognitive domains, such as episodic memory, working memory, fluid intelligence, and perceptual speed. An overview of all assessed tasks in the acquired order can be gained from Table 1. Many of the tasks have been used and described in detail before (Schmiedek, Bauer, et al., 2010; Schmiedek, Lövdén, et al., 2010). As the tasks measuring fluid intelligence were a somewhat last-minute addition, they were placed directly after each other at the very end of our cognitive testing session. In hindsight, we believe this was not ideal and it might have been better to split those cognitive tests into two sessions altogether, to lower cognitive demand and thereby potentially heighten motivation.

**Table 1.** Overview of all cognitive tasks in order administered

| Cognitive Domain | Task | Approximate duration |
| --- | --- | --- |
| Episodic memory | Item-Context task | 20 minutes |
| Working memory | Spatial Updating | 20 minutes |
| Episodic memory | Verbal Learning and Memory Test (VLMT) | 20 minutes |
| Perceptual speed | Digit Symbol Substitution task | 5 minutes |
|  | *BREAK* | 10 minutes |
| Episodic memory | Repeat VLMT | 5 minutes |
| Episodic memory | Rey-Osterrieth Complex Figure test (Rey Figure) | 5 minutes |
| Vocabulary | Word recognition | 5 minutes |
| Episodic memory | Repeat Rey Figure | 5 minutes |
| Working memory | Number N-Back | 15 minutes |
| Working memory | Letter Updating | 15 minutes |
| Episodic memory | Repeat Rey Figure | 5 minutes |
|  | *BREAK* | 20 minutes |
| Perceptual speed | Figural Speed | 10 minutes |
| Perceptual speed | Numerical Speed | 10 minutes |
| Perceptual speed | Verbal Speed | 10 minutes |
| Episodic memory | Face-Profession task | 20 minutes |
|  | *BREAK* | 10 minutes |
| Episodic memory | Object Location task | 20 minutes |
| Fluid intelligence | Practical Problems | 10 minutes |
| Fluid intelligence | Figural Analogies | 10 minutes |
| Fluid intelligence | Letter Series | 10 minutes |

##### Episodic memory

###### Item-Context Task

The Item-Context task was adapted from a task described in (Berron et al., 2018). Participants were presented stimuli (computer-generated with 3ds Max, Autodesk Inc., San Rafael, USA; isoluminant) comprising everyday indoor objects presented on a grey background or empty indoor scenes (i.e., rooms). Lure stimuli were created by changing certain spatial features of objects (shape, but not color, position, or size) or room (geometry, but not color or viewpoint). During a trial, four stimuli were presented on a touchscreen for 3 seconds each; the first two stimuli were always new images and were either both objects or both rooms, the third stimulus was either a repeat or the lure of the first stimulus, and the fourth was either a repeat or the lure of the second stimulus. During presentation of the third and fourth stimuli, participants were instructed to use a stylus pen to either touch a field on the bottom of the screen to indicate if they had already seen the exact same image (repeat), or touch the area of the picture they believed had changed (lure). The presentation of object vs. room and repeat vs. lure was randomized and counterbalanced. In total, participants were presented 56 four-stimulus trials for a total of 224 stimuli (56 object first presentations, 28 object repeat, 28 object lure, 56 room first presentations, 28 room repeat, 28 room lure). Corrected hit rate was calculated as hit rate (percent correct repeat items) – false alarm rate (percent incorrect lure items). Corrected hit rate is considered an unbiased measure of old-new discrimination used to assess recognition memory (Snodgrass & Corwin, 1988).

###### The Verbal Learning and Memory Test

(VLMT) (Helmstaedter & Durwen, 1990) is a German version of the Rey Auditory Verbal Learning Test. Participants heard a list of 15 nouns, serially presented via headphones. Presentation of the list was followed by a recall phase in which a computer screen prompted the participants to type as many words as they could remember from the list. This was repeated over five learning trials, each with the same word list. After that, an interference list was presented. Yet again, free recall was tested directly after the interference list as well as 30 min later (late recall), followed by a final recognition test. The same list of German nouns was used for all participants. The sum of items recalled across trials 1 to 5 provides a measure of overall learning performance and will be used here further on as the manifest variable.

###### The Face–Profession task

assesses associative binding on the basis of recognition of incidentally encoded face–profession pairs. During the study phase, 45 face–profession pairs were each presented for 3.5 s on the computer screen and the participants had to indicate via button presses whether the faces matched the profession or not. After a 3-min delay between study and test phase, 54 face–profession pairs consisting of 27 old pairs, 9 new pairs, and 18 newly arranged pairs were presented (in newly arranged pairs the shown face is the same, but is associated with a new profession). Participants were asked to decide whether they had seen a given face–profession combination before or not and to rate the confidence of their decision on a three-point scale ranging from 1 = not sure to 3 = very sure.

###### In the Object Location task

sequences of 12 colored photographs of real-world objects were displayed at different locations in a 6-by-6 grid. After presentation, objects appeared on the side of the screen and had to be moved to the correct locations by clicking on the objects and the locations with the computer mouse. There was one practice trial and two test trials.

###### The Rey-Osterrieth Complex Figure

(Rey, 1941). The complex figure test consisted of 18 different geometric patterns. First, the figure had to be copied in as detailed a manner as possible (copy trial). After a 3 min (early recall) and a 30 min (late recall) delay interval, participants were asked to reproduce the figure as accurately as possible from memory. This was followed by a recognition memory test that consisted of 12 single elements of the 18 scoring elements of the original figure along with 12 new elements from the alternative figure (i.e. from the Modified Taylor Figure) serving as lures. Participants were instructed to circle the elements that were part of the original design that had been copied and drawn. The recognition memory test of the complex figure test required discrimination between similar looking complex objects and thus posed high demands on accurate memory representations and pattern separation.

##### Working memory

###### In the Letter Updating task

participants saw sequences of the letters A, B, C, and D. Each sequence contained seven, nine, eleven, or thirteen letters. After each sequence, participants had to report the last three letters of the sequence in the same order in which they had appeared via buttons corresponding to each of the four letters.

###### In the Spatial Updating task

participants were asked to remember and update the positions of dots within 3 × 3 grids. Two or three grids were displayed simultaneously for 4000 ms, each with a blue dot in one of the nine possible positions. Arrows were then presented for 2500 ms below each grid, and participants were instructed to mentally shift each dot according to the corresponding arrow. After each dot had been shifted twice, participants indicated the resulting end position of each dot via mouse click.

###### In the Number-N-Back task

single digits (1–9) were presented successively in three visible boxes, and participants had to decide whether the currently presented integer matched the one shown three presentations earlier, i.e., previously presented in the same box.

Responses were indicated via button press with left and right index fingers.

##### Fluid intelligence (Gf)

###### The Practical Problem task

consisted of 12 items depicting everyday problems such as the times in a bus schedule, instructions for medication, a warranty for a technical appliance, a rail map, as well as other forms and tables. For each item, the problems were presented in the upper part of the screen, and five alternative responses were shown in the lower part. Subjects responded by clicking on one of the five alternatives with the computer mouse. A single practice item was provided. The test phase terminated when subjects had made three consecutive errors, or had reached the maximum time limit of 10 minutes, or had reached the last test item. Items were ordered by difficulty.

###### Figural Analogies

Items in this test followed the format “A is to B as C is to ?”. One figure pair was presented in the upper left part of the screen and a single figure was shown beside it. Participants had to use the same rule as the one applying to the complete figure pair to choose one of the five alternative responses presented below. Subjects entered their response by clicking on one of the five alternatives with the mouse. Before the test phase, instructions and three practice items were given. The test phase terminated when subjects had made three consecutive errors, had reached the maximum time limit (10 min), or had reached the last test item. Items were ordered by difficulty.

###### The Letter Series task

consisted of 22 items. Each item contained five letters followed by a question mark (e.g., c e g i k ?). Items were displayed in the upper half of the screen, and five response alternatives were presented in the lower half. Items followed simple rules such as +1, –1, +2, or +2 +1. Subjects entered their response by touching one of the five answer alternatives. The score was based on the total number of correct responses. Instructions and three practice items were given before the test phase. Again, the test phase terminated when subjects had made three consecutive false responses, had reached the maximum time limit (6 min), or had reached the last item of the test. Items were ordered by difficulty.

##### Perceptual speed (numerical, verbal, and figural)

Perceptual speed was evaluated with three comparison tasks, in which participants had to quickly decide if the two stimuli being presented were the same or different, and respond with a button press with either the left or right hand, respectively. For the numerical version, two strings of four digits (1–9) were presented on the left- and right-hand sides of the screen. For the verbal version, two strings of four letters were presented. For the figural version, objects called “fribbles” were presented. Fribbles are three-dimensional colored objects comprising several connected components (images courtesy of Michael J. Tarr, Carnegie Mellon University, http://tarrlabwiki.cnbc.cmu.edu/index.php/Tarr). Stimuli were either identical or differed on a single aspect (i.e., one digit, letter, or component).

*Digit Symbol test* (DS) (Wechsler, 1981). The DS test consisted of a code box with nine digit–symbol pairs, where each digit was paired with a corresponding symbol, and rows of double boxes, each with a digit in the top box and an empty lower box. Participants were asked to fill in as many corresponding symbols as possible in 90 s. The final score indicates the number of correctly filled boxes, with penalty for wrong answers (score correct – wrong).

##### Spot-a-word test (MWT-A)

The spot-a-word test (Lehrl et al., 1991) is a paper-pencil multiple choice vocabulary test, assumed to also be an indicator for crystalline intelligence. It consisted of 37 word lines, whereby participants had to select in each line one real word out of four meaningless words.

#### Psychosocial functioning assessment

A set of self–report questionnaires that have been validated and applied already within the BASE–II studies (Gerstorf et al., 2015) were administered to explore potential correlates of individual differences in the psychosocial assessment to the different training-regimes. This psychosocial assessment contained measures of self-reported health, well-being, stress, coping, and attitudes towards aging. After the first cognitive assessment, participants were asked to fill out a comprehensive psychosocial take-home questionnaire and return this during their next visit at the institute. The same procedure was applied for the post psychosocial questionnaire.

First, demographics and person characteristics such as height, weight, marital status (1*–*single, 2*–*widowed, 3*–*divorced, 4*–*married, 5*–*partnership), and highest school graduation (1*–*elementary school, 2*–*secondary school, 3*–*secondary education, 4*–*final secondary school qualification, 5*–*graduate degree) were asked. The next set of questions pertained to medication and health status. Participants had to indicate which and how many medications they take regularly. Participants had to check whether they had been diagnosed by a doctor with one or more of the following diseases: diabetes, heart disease, stroke, high blood pressure, depression, joint disease, chronic back pain, cancer. If so, they had to indicate which year the diagnosis has been made.

##### Obligatory and optional personal life investment (PLI)

(Schindler & Staudinger, 2008). PLI was assessed as the amount of energy invested in terms of action and thought in ten life domains (one item per domain): health, cognitive fitness (cognition), hobbies and interests (leisure), friends and acquaintances (friends), sexuality, well-being of family members (family), occupation or occupation-like activities (occupation), independence, thinking about one’s life (life reflection), and one’s death and dying (death). For each life domain, participants were asked to respond on a 5–point Likert Scale ranging from “never” to “very often”: “How about your health? How much do you presently think about it or do something about it?”. Next, all 10 life domains were listed and participants had to check the one most important current life domain they are currently thinking about or doing the most for. The last set of questions referred to participants’ life when they were between 30 and 50 years old. Participants were instructed to remember how the importance of life domains had changed compared to the age between 30-50 years. A different 5–point Likert Scale was used here ranging from “less than before” to “more than before”.

##### Life satisfaction

was assessed with one item: “How satisfied are you currently all in all with your life?”. Participants rated their satisfaction with their current life circumstances on a scale ranging from 0–very unsatisfied to 10–very satisfied.

##### Well–being

We used the Satisfaction with Life Scale to assess the cognitive– evaluative component of well–being (Diener et al., 1985) (Cronbach’s alpha .88) in which participants rated five statements (e.g., “In most ways my life is close to my ideal”) on a 5– point scale ranging from 1 (strongly disagree) to 5 (strongly agree) to indicate their level of agreement.

##### Perceived stress

We used an 8–item version of the Perceived Stress Scale (PSS) to assess perceived stress within the last four weeks (Cohen et al., 1994). Participants rated eight statements (e.g., in the last month, how often have you felt nervous and “stressed”?) on a 5– point Likert Scale ranging from 0 (never) to 4 (very often) to indicate their level of agreement.

##### Future time perspective scale

We used the 10–item Future Time Perspective scale (Carstensen & Lang, 1996). Participants responded to each item (e.g., “Many opportunities await me in the future”.) using a Likert scale ranging from 1 (very untrue for me) to 7 (very true for me). The scale composite consists of the unit–weighted mean across items, with lower scores indicating a more limited general future time horizon (Kozik et al., 2020).

##### Internal and external control beliefs

The four control belief dimensions were assessed using items derived from conceptual and empirical work on locus of control (Drewelies et al., 2021; Kunzmann et al., 2002). For each dimension, participants were asked to indicate the extent to which they agreed with three or four statements using a 5–point Likert Scale from 1–does not apply to me at all to 5–applies very well to me. Sample items include “I can make sure that good things come my way.” (internal control beliefs over desirable outcome); “It’s my fault if something goes wrong in my life.” (internal control beliefs over undesirable outcomes); “The good things in my life are determined by other people.” (external control beliefs in powerful others); and “The good things in my life are, for the most part, a matter of luck.” (external control beliefs in chance).

##### Goal adjustment capacities

were assessed using the 10-item Goal Adjustment Scale (Wrosch et al., 2003). The scale measured individual differences in peoples’ tendencies to disengage from unattainable goals (goal disengagement) and to re-engage with new alternative goals (goal reengagement). Participants were asked to indicate how they usually reacted to situations where they were forced to stop pursuing important goals using 10–point scales (0–strongly disagree, 10–strongly agree). Four items assessed goal disengagement (e.g., “It’s easy for me to reduce my effort towards the goal”), and six items assessed goal reengagement (e.g., “I seek other meaningful goals”).

##### Subjective age

Perceived age was assessed with the item “How old do you feel?”. Participants were asked to write down a specific age that they felt.

##### Social networks ag

Participants had to rate the average age of their friends within their social network.

##### Loneliness

Loneliness was measured utilizing the UCLA Loneliness Scale (Russell, 1996) consisting of seven items. Participants were asked to rate each statement on a 5–point Likert Scale ranging from 1–5 (‘1–strongly disagree’ to ‘5–strongly agree’). Higher scores indicate stronger feelings of loneliness (Cronbach’s α = .81; Drewelies et al., 2021).

##### Assessment of subjective memory impairment

The following items concerning subjective memory complaints were taken from (Jessen et al., 2007): “Do you feel like your memory is becoming worse?”. Possible answers were “yes”, “no”, and “If yes, since when?”. “Do you think these changes are normal for your age group?”. Possible answers were “yes” and “no”, and “Are you concerned about these changes?”.

Additionally, participants were asked the following questions (1) “Have you ever consulted a doctor due to memory problems?” including possible answers: “yes”, “no”, “if yes, when?” (2) “How often do you say something like: “I can’t remember”, or “I can’t remember anything”, or “I forget too much”?” Possible responses were the following (1) daily (2) weekly (3) monthly, and (4) never.

##### The Subjective Health Horizon Questionnaire (SHH-Q)

The SHH-Q is a validated new self-report measure (SHH-Q) (Düzel et al., 2016) with 30 items that assesses s individuals’ future time perspectives in relation to four interrelated but distinct lifestyle dimensions: novelty-oriented exploration (Novelty), bodily fitness (Body), work goals (Work), and goals in life (Life Goals). Each SHH-Q item was accompanied by a vector representation of the lifespan underneath, ranging from 10 years of age on the left to 110 years of age. Participants were asked to respond to each item by indicating first their current chronological age and then the maximum future age at which they estimate they would mentally and physically still be able to perform the activity in question.

##### EPIC Questionnaire

We administered the European Prospective Investigation into Cancer and Nutrition (EPIC) food frequency questionnaire (FFQ) to assess daily average dietary tyrosine intake in grams (Boeing et al., 1997). The questionnaire assesses dietary intake of 148 foods during the last 12 months. For each item and given portion size (indicated by means of line drawings), participants rated the frequency of consumption. Then, we derived the nutrient concentrations for each food item of the FFQ by taking values provided by the Federal Coding System.

##### The Leisure Activity Questionnaire

was designed to assess the number of hours per week occupied by 18 diverse cognitive activities (e.g. driving the car, playing computer games, cooking, reading books or newspapers, attending university lectures, playing cards, instruments, or writing letters), 18 physical activities (e.g. gardening, cleaning, walking, cycling, dancing, jogging, hunting, stretching), and 18 social activities (e.g. spent time with family, meeting friends, phone calls with relatives, visiting a pub or restaurant) on a scale ranging from zero hours to 15+ hours per week. Next, participants had to indicate on a 5– point Likert Scale how cognitively challenging they rated these activities, ranging from “easy” to “hard”.

##### Reading preferences and behaviour

This group-specific questionnaire was administered within the active control group only. Reading behavior was assessed with seven items (“I like to read stories.”). Responses were recorded using a 6-point scale (1–does apply to 6– does not apply). Eight items about one’s own thoughts and attitudes were assessed using a 10-point scale (1–strongly disagree, 10–strongly agree).

##### Spanish proficiency test score

The unforeseen test was administered in the LG and L+EG only, in the last group session under the supervision of the Spanish teacher. It was prepared by the Spanish teacher based on the material covered in the language learning application Babbel. It required translating a total of 55 items, either words (e.g. ‘school’), common phrases (e.g. ‘Nice to meet you’), sentences (e.g. ‘I like to dance’) or questions (e.g. ‘How is the weather’) from German to Spanish. The maximum attainable score was 110.

#### Training data collection

During the six-month training period, participants performed their respective training paradigms at home. Therefore, data was also collected at home on an almost-daily basis via tablets, ergometers, and a fitness tracking device, to document training progress. At the beginning of each at-home training session using the provided tablets (and bicycle ergometers, in the case of EG and L+EG), participants were asked to respond to questions on the tablet assessing their current emotional state (all groups; How happy are you feeling today? How stressed are you feeling today? Do you have the feeling you can influence what is happening in your life right now?) as well as motivational state (LG, EG, and L+EG; How well do you think you will train today? How motivated do you feel today?) on visual analog scales ranging from 0–100.

Language-learning related parameters were collected within the Babbel application and included number of overall items done per session, number of items correct, time spent with the app. Babbel is a subscription-based language learning app, it’s original learning content is developed by the company’s team of educators and linguists. There are beginner, intermediate and grammar courses, vocabulary lessons, as well as courses with tongue-twisters, idioms, colloquialisms, and sayings. Participants could choose between different courses to keep them engaged and the language learning always interesting.

Exercise-related parameters were collected from participants who exercised during ergometer training (in the EG and the L+EG) through the ergometer and its handlebars with the data being relayed to the tablet via Bluetooth. These parameters included time spent training, mean pulse during training, and mean ergometer resistance during training. The ergometer training app was developed by the neotiv team (www.neotiv.com).

As for reading behavior, reading time was recorded of all participants, from within the reading app. Participants received their reading material peu à peu directly on their tablets. The texts ranged from short stories (for example, by Alice Munro, James Salter, Judith Hermann) to novels (for example, A Handmaid’s Tale by Margaret Atwood, Sherlock Holmes by Sir Arthur Conan Doyle), all in German.

Additionally, 160 participants across the four groups, also including some participants that were drop-outs eventually, consented to wearing an activity tracking bracelet (www.fitbit.com) for the duration of the study, also while they slept. Acquired activity tracker parameters included kilocalories, steps, and distance walked. During sleep, measures of time asleep, time spent in bed, times waking up, and periods and durations of restlessness were recorded. Note, however, that actual wear-time differed greatly between participants.

## Conclusion

Around the world, and especially in Europe and the US, the older segment of the adult population is growing in size and proportion (Vaupel et al., 2003). In dealing with the demographic change, it is important what people make of the added years of their lives. Cognitive performance is known to decrease with increasing age, even in healthy individuals. This decline starts as early as around 30 years for some cognitive abilities, and accelerates after about 65 years of age. Decline of cognitive functioning is a health risk, and it is also experienced as such. Declining cognition also puts a burden on the health-care system and the labor market. Preventing or attenuating cognitive decline is probably the most effective measure to postpone the age at which individuals are no longer able to lead an independent life.

Effective pharmacological treatments of cognitive decline as well as dementia are currently still unavailable. Some modifiable lifestyle factors, such as poor diet and physical and cognitive inactivity have, however, been identified to be associated with the risk for cognitive decline, not only in neurodegenerative diseases but also in normal aging (Lindenberger, 2014). Furthermore, physical exercise can convey a protective effect, even if initiated in advanced old age (Tolppanen et al., 2015) and challenging cognitive activities have been shown to improve performance in trained tasks (Brehmer et al., 2007; Schmiedek, Lövdén, et al., 2010) and decelerate age-related structural decline (Lövdén et al., 2012; Wenger et al., 2012). Demonstrating positive effects of a novel form of intervention that combines physical exercise with cognitive stimulation would pave the way for fostering lifestyles that extend quality of life and functional independence.

We implemented an easily accessible and inherently motivating at-home training program that is expected to promote successful aging in older adults.Training paradigms were deliberately embedded in participants’ daily lives to enable better generalization of the knowledge gained. With our study design, we can investigate the effects of language learning, physical exercise training on stationary bicycles and a combination of both on healthy older adults on the neural, cognitive, and affective level.

## Acknowledgements

This work was supported by the Max Planck Society and the Max Planck Institute for Human Development and is part of the BMBF funded EnergI Consortium (01GQ1421B). We are very grateful to the Babbel team for providing us with free access to their language learning app, and to the Neotiv team for providing the app for ergometer training as well as their technical support. We thank Maria Bautista-Sanchez and Matthew Youlden as our Spanish instructors, Katrin Sohnrey as our toning and stretching instructor, and Carsten Sommerfeldt and everyone at Shared Reading for organizing our book club. We thank Kirsten Becker, Anke Scheepers-Klingebiel and Ludmila Müller for invaluable organizational assistance, Sebastian Schröder and his team for their help with all technical equipment, the MRI team at the Max Planck Institute for Human Development (Sonali Beckmann, Nadine Taube, Thomas Feg, and Davide Satoro) and all participants for their time and support, as well as Maike Kleemeyer for valuable comments on an earlier draft of this manuscript.

## Footnotes

1 Approximately half of the final sample investigated here was also part of the longitudinal Berlin Aging study II (BASE-II), offering a rich set of additional variables such as socioeconomic background, extensive medical and genetic data.

2 The original attempted age range of the study was 65 to 75 years. Because of interested spouses of volunteers, we released this inclusion criterion slightly and in the end included participants aged 63 to 78 years old.

